# Hierarchical Model for the Role of J-Domain Proteins in Distinct Cellular Functions

**DOI:** 10.1101/2020.06.12.147645

**Authors:** Shinya Sugimoto, Kunitoshi Yamanaka, Tatsuya Niwa, Yuki Kinjo, Yoshimitsu Mizunoe, Teru Ogura

## Abstract

In *Escherichia coli*, the major bacterial Hsp70 system consists of DnaK, three J-domain proteins (JDPs: DnaJ, CbpA, and DjlA), and one nucleotide exchange factor (NEF: GrpE). JDPs determine substrate specificity for the Hsp70 system; however, knowledge on their specific role in bacterial cellular functions is limited. In this study, we demonstrated the role of JDPs in bacterial survival during heat stress and the DnaK-regulated formation of curli—extracellular amyloid fibers involved in *E. coli* biofilm formation. Genetic analysis with a complete set of JDP-null mutant strains demonstrated that only DnaJ is essential for survival at high temperature, while DnaJ and CbpA are indispensable in DnaK regulation of curli production. Additionally, we found that DnaJ and CbpA are involved in the expression of the master regulator CsgD through the folding of MlrA; this keeps CsgA in a translocation-competent state by preventing its aggregation in the cytoplasm. Our findings support a hierarchical model wherein the role of JDPs in the Hsp70 system differs according to individual cellular functions.

## INTRODUCTION

Proteostasis is the maintenance of protein homeostasis in cells and is essential for all life. Key players in proteostasis include molecular chaperones, which assist in protein folding, refolding of denatured and aggregated proteins, protein transport, and quality control of regulatory proteins. Therefore, molecular chaperones are involved in diverse cellular activities including cell division, DNA replication, stress response, organelle functions, and autophagy (Hipp et al., 2019).

The 70-kDa heat shock proteins (Hsp70s) are ubiquitous molecular chaperones involved in a wide variety of cellular functions (Mayer and Kityk, 2015). Hsp70s are ATP-dependent molecular chaperones that consist of an N-terminal nucleotide-binding domain (NBD) and a 25 kDa C-terminal substrate-binding domain (SBD) (Zhu et al., 1996). Hsp70s function via nucleotide-regulated substrate binding and release cycles (Szabo et al., 1994; McCarty et al., 1995). In the ATP-bound state, Hsp70s exhibit low affinity toward substrates; therefore, their rates of substrate binding and release are rapid. In contrast, the ADP-bound state exhibits high substrate affinity, with consequent low rates of substrate binding and release. In the nucleotide-dependent chaperone cycle, Hsp70s require cofactors, known as co-chaperones. Among co-chaperones, J-domain proteins (JDPs), also referred to as Hsp40s, stimulate the ATPase activity of Hsp70s and the binding of substrate proteins (Gässler et al., 1998; Suh et al., 1998, 1999). Conversely, nucleotide exchange factors (NEFs), another type of co-chaperone, induce ADP dissociation from the NBD of Hsp70s and substrate-release from the SBD (Brehmer et al., 2004). Although the amplification and diversification of Hsp70s may be involved in their functional versatility, JDPs far outnumber Hsp70s in the vast majority of life forms (Kampinga and Craig, 2010), and the multiplicity of JDPs drives the functional diversity of Hsp70s (Craig and Marszalek, 2017). Six JDPs (DnaJ, CbpA, DjlA, HscB, DjlB, and DjlC) have been identified in *Escherichia coli*, 22 in *Saccharomyces cerevisiae*, and 41 in humans, whereas there are three Hsp70s in *E. coli* (DnaK, HscA, and HscC), 16 in *S. cerevisiae*, and 17 in humans (Table EV1) (Powers and Balch, 2013). In *E. coli*, three JDPs (DnaJ, CbpA, and DjlA) productively interact with DnaK and play redundant roles in the regulation of DnaK chaperone activity (Sell et al., 1990; Ueguchi et al., 1994; Genevaux et al., 2001; Gur et al., 2004). These observations suggest that DnaJ, CbpA, and DjlA share overlapping functions.

Previously, we demonstrated that DnaK (a bacterial Hsp70) serves an important role in the formation of *E. coli* biofilms—well-organized microbial communities that form on surfaces—and that the production of curli—extracellular functional amyloid fibers—relies on DnaK functions (Arita-Morioka et al., 2015). Curli fibers play crucial roles in biofilm organization and host colonization by adhering to surfaces and holding bacterial cells in a self-produced extracellular matrix (Olsén et al., 1989; Chapman et al., 2002). Secretion and assembly of curli is mediated by a characteristic secretion pathway, known as the nucleation-precipitation mechanism or the type VIII secretion system (Desvaux et al., 2009). In *E. coli*, seven proteins encoded by two dedicated operons, the curli-specific genes *BAC* (*csgBAC*) and *DEFG (csgDEFG)* operons, are involved in the expression, export, and assembly of the amyloid fibers (Hammar et al., 1995). An alternative sigma factor, RpoS, also known as σ^S^ or σ^38^, activates expression of the *csgDEFG* operon (Hammar et al., 1995; Dudin et al., 2014). In addition, CsgD, the master transcriptional regulator of curli synthesis, directly promotes transcription of the *csgBAC* operon (Hammar et al., 1995; Zakikhany et al., 2010). CsgA and CsgB are the major and the minor curli subunits, respectively. Following transport across the cytoplasmic membrane via the Sec translocon, the CsgA and CsgB subunits are exported across the outer membrane in a manner dependent on CsgG, a curli-specific translocation channel (Goyal et al., 2014; Cao et al., 2014). After secretion, CsgB nucleates CsgA subunits into amyloid fibrils (Shu et al., 2012). In addition, CsgE, a periplasmic accessary protein, directs CsgA to CsgG for secretion (Nenninger et al., 2011). CsgF, an extracellular accessory protein, is required for the specific localization and/or nucleation activity of CsgB (Nenninger et al., 2009).

Recently, we found that DnaK multitasks to maintain homeostasis of the key players in curli biogenesis, including regulation of the quantity and *de novo* folding of RpoS and CsgD, and export of CsgA (Sugimoto et al., 2018); however, curli production was not affected by single knockout of JDPs (DnaJ, CbpA, and DjlA), all of which are known to functionally interact and cooperate with DnaK (Genevaux et al., 2001, 2007). These findings motivated us to investigate whether DnaK works alone or together with specific JDPs in distinct cellular functions, such as curli biogenesis and survival at high temperature. Our results will further our understanding of the role of JDPs and the activity of the DnaK chaperone system.

## RESULTS

### Either DnaJ or CbpA is indispensable for curli biogenesis

To address which JDPs are essential for curli biogenesis, we used the Keio collection, a widely used *E. coli* single-gene knockout library (Baba et al., 2006). We also constructed a complete set of JDP double- and triple-null mutants of the K-12 strain BW25113 (Table EV2) by the one-step method for inactivation of chromosomal genes (Datsenko and Wanner, 2000). Curli production was detected on Congo Red-containing YESCA agar (CR-YESCA: 1% casamino acids, 0.1% yeast extract, and 2% agar) plates at 25°C for 2 days. In the strains Δ*cbpA* Δ*dnaJ* and Δ*cbpA* Δ*djlA* Δ*dnaJ*, curli production was reduced, as well as in the strains Δ*dnaK*, Δ*csgA*, Δ*csgD*, Δ*csgG*, and Δ*rpoS*, while it was not affected in the others (Fig 1A). Curli was also evaluated by immunoblotting for CsgA monomers as described below.

**Figure 1.**
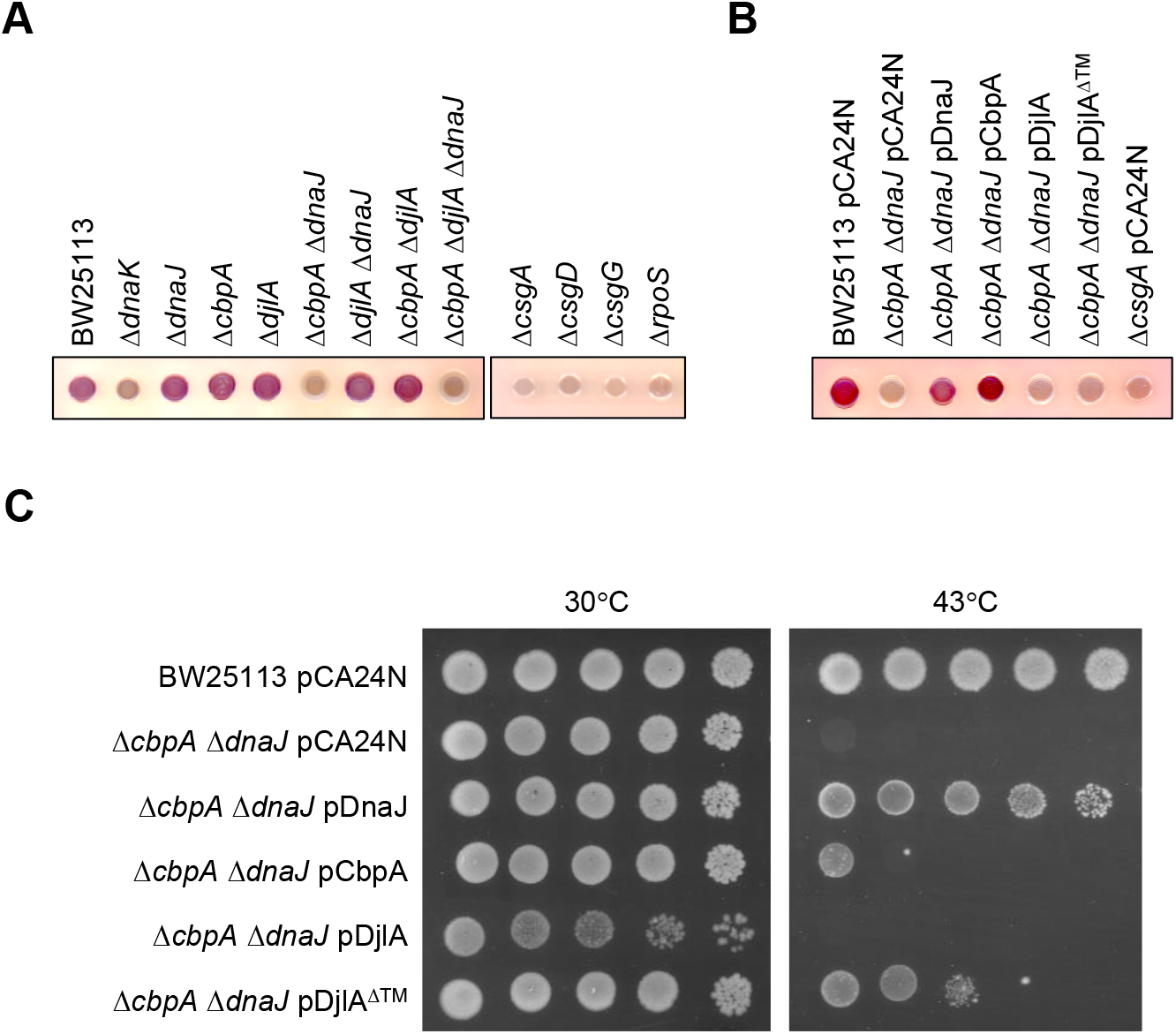
Effects of JDP deletion on curli biogenesis. (A) Curli production in *E. coli* BW25113 and its isogenic mutants were examined on Congo Red (CR)-containing YESCA plates (CR-YESCA). Strains were grown at 25°C for 2 days. The strains Δ*csgA, ΔcsgD, ΔcsgD*, and Δ*rpoS* were used as negative controls. (B) Curli production in BW25113 and its isogenic Δ*cbpA* Δ*dnaJ* strains transformed with pCA24N (empty vector) and the indicated JDP-expression plasmids were examined on CR-YESCA plates supplemented with chloramphenicol. Strains were grown at 25°C for 2 days. (C) Thermosensitivity of the indicated strains was assayed on LB agar plates supplemented with chloramphenicol. Ten-fold serial dilutions of the overnight cultures were spotted on the plates. Plates were incubated at 30°C or 43°C for 24 h.

To confirm the responsibility of JDPs in the observed phenotypic changes, we conducted a trans-complementation assay using the JDP-expression plasmids (Table EV2). The plasmids carrying DnaJ and CbpA restored curli production in BW25113 derivatives Δ*cbpA* Δ*dnaJ* (Fig 1B). In contrast, neither DjlA nor DjlA^ΔTM^, which lack the transmembrane domain, restored curli production. These results indicate that either DnaJ or CbpA, but not DjlA, is essential for curli production and that DnaJ and CbpA work redundantly in this process.

### DnaJ is essential for survival at high temperature

Previously, the requirement for JDPs in the survival of *E. coli* at high temperature was reported using MC4100 and its isogenic mutants (Sell et al., 1990; Ueguchi et al., 1994; Genevaux et al., 2001). These reports showed that deletion of JDPs was not lethal at 30°C and that DnaJ, but neither CbpA nor DjlA, was essential for the growth of MC4100 at 43°C. We revisited the requirement of JDPs in the survival of BW25113 at high temperature using newly constructed null mutant strains (Table EV2). Our study also revealed that all JDPs were dispensable for growth of BW25113 at 30°C, and only DnaJ was indispensable for survival at 43°C (Fig EV1A). In addition, complementation analysis demonstrated that expression of DnaJ rescued the growth of BW25113 Δ*dnaJ* Δ*cbpA* as well as BW25113 Δ*dnaJ* Δ*cbpA* Δ*djlA* at 43°C (Fig 1C and EV1B). Expression of CbpA or DjlA^ΔTM^ partially recovered the survival of BW25113 Δ*dnaJ* Δ*cbpA* and Δ*dnaJ* Δ*cbpA* Δ*djlA* at 43°C (Fig 1C and EV1B). These results indicate that DnaJ plays the most pivotal role in survival of *E. coli* at high temperature.

### Diversity of DnaK chaperone activities required for distinct phenotypes

Our results indicate that under physiological conditions, only DnaJ is essential for the growth of *E. coli* at high temperature, while DnaJ and CbpA work redundantly in curli production. These findings imply the presence of a diversity of DnaK chaperone activities required for different cellular processes. If this is the case, some DnaK mutants with reduced chaperone activity would support curli production but not in survival at high temperature. Here, we focused on a mutant of DnaK with reduced interaction with JDPs. Substitutions of Tyr-145, Asn-147, and Asp-148 to Ala in DnaK (DnaK^YND^) (Fig 2A) caused reduced interaction with DnaJ via its J-domain (Gässler et al., 1998), suggesting that this mutant DnaK interacts very weakly with JDPs. We also focused on defective allosteric communication between the NBD and SBD. Previously, two charged residues, a surface-exposed, positively charged residue in the NBD (Lys-155) and a negatively charged residue in the linker connecting NBD and SBD domains (Asp-393), were shown to be important for interdomain communication (Vogel et al., 2006). One of the mutants, DnaK^K155D^, in which Lys-155 in NBD is replaced with Asp, greatly reduces the stimulation of the substrate dissociation rate by ATP while retaining substrate- and DnaJ-mediated stimulation of ATPase activity (Vogel et al., 2006). In contrast, another mutant, DnaK^D393A^, in which Asp-393 is substituted to Ala, drastically reduces the stimulation of substrate dissociation rate by ATP, as well as the substrate- and DnaJ-mediated stimulation of ATPase activity (Vogel et al., 2006).

**Figure 2.**
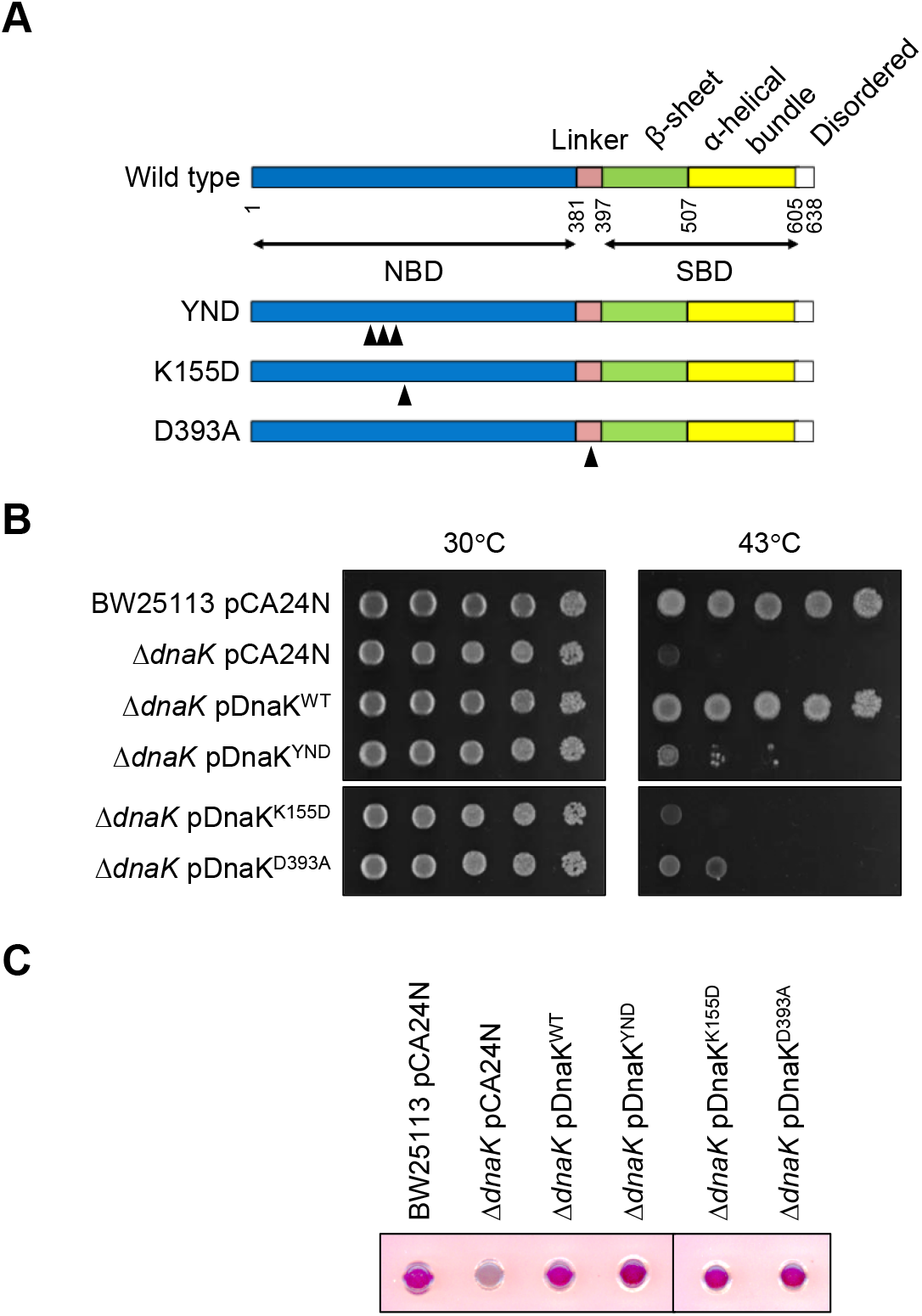
Complementation of the *dnaK-null* strain with defective DnaK mutants in cooperation with JDPs and interdomain communication. (A) The domain structure of DnaK and mutants used in this study. Arrowheads represent mutation sites. (B) The thermosensitivity of the indicated strains was assayed on LB agar plates supplemented with chloramphenicol. Ten-fold serial dilutions of the overnight cultures were spotted on the plates. Plates were incubated at 30°C or 43°C for 24 h. (C) Curli production in the strains was examined on CR-YESCA plates supplemented with chloramphenicol. The plates were incubated at 25°C for 2 days.

In our study, expression of these mutant DnaK proteins did not induce recovery from the growth defect of the Δ*dnaK* strain at high temperature (Fig 2A and B). This indicates that interaction of DnaK with JDPs and allosteric communication between the NBD and SBD of DnaK is indispensable for survival under heat stress conditions, as previously reported (Gässler et al., 1998; Vogel et al., 2006). However, notably, expression of DnaK^YND^, DnaK^K155D^, and DnaK^D393A^ fully restored curli production in the Δ*dnaK* strain (Fig 2C).

These results support our hypothesis that there is a functional diversity of the DnaK system required for distinct cellular functions.

### Loss of DnaJ and CbpA reduces expression of CsgD

Based on our data, we hypothesized that either DnaJ or CbpA plays an important role for expression and folding of certain proteins associated with curli biogenesis. To address this, we investigated the expression levels of several proteins in the BW25113 strains by immunoblotting.

Firstly, we confirmed the expression of chaperone proteins (Fig 3A). As expected, DnaK, DnaJ, and CbpA were not detected in the respective mutant strains, revealing that deletion of these proteins was successfully conducted. Only DjlA was not detected because of the absence of an available specific antibody. The cellular level of DnaK was slightly increased in the strains lacking DnaJ, while that of DnaJ was much higher in the strain Δ*dnaK* compared to the other strains. These phenomena can be accounted by the accumulation of RpoH, also known as σ^32^ or σ^H^, in these mutant strains, which induces expression of DnaK and DnaJ (Tatsuta et al., 2000). These results are also consistent with the previous observation (Gur et al., 2004). In addition, the cellular level of CbpA in the strain Δ*rpoS* was lower than that in the wild-type strain, confirming that the expression of CbpA is positively regulated by RpoS during the stationary growth phase as previously reported (Yamashino et al., 1994).

**Figure 3.**
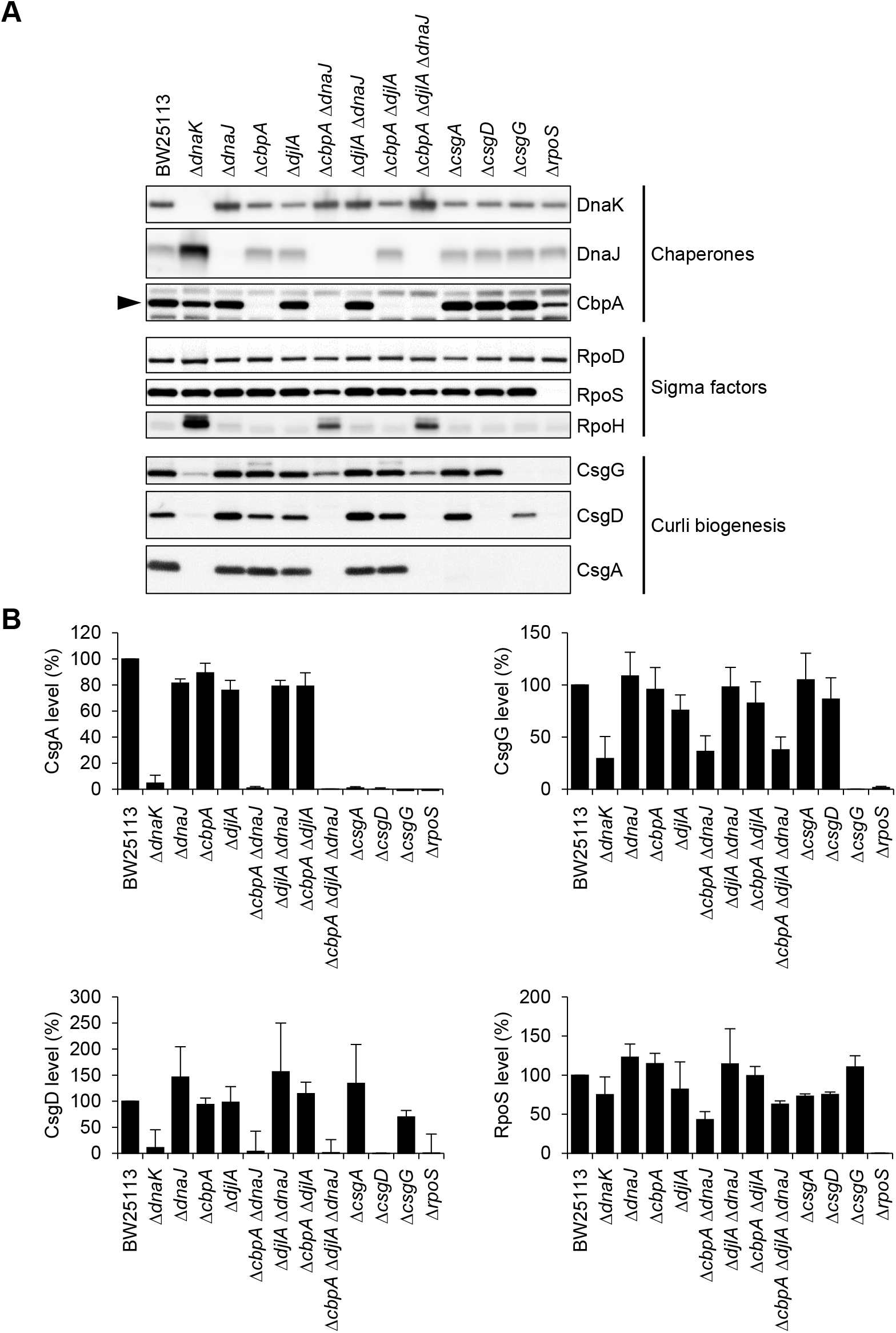
Effects of JDPs on expression of curli-related proteins. (A) BW25113 and its isogenic null mutants were grown on YESCA plates at 25°C for 2 days. Expression of curli-related proteins, chaperones, and sigma factors was analyzed by immunoblotting. CsgA monomers were depolymerized with hexafluoroisopropanol (HFIP). All experiments were conducted using total protein samples. Relative protein levels of CsgA, CsgD, CsgG, and RpoS were quantified based on the band intensity of immunoblots. All experiments were repeated at least three times to ensure accuracy and averaged values with standard deviations were calculated. The band intensities in the BW25113 parental strain were defined as 100%.

Secondly, to elucidate the mechanisms of how either DnaJ or CbpA affects the biogenesis of curli, we investigated cellular levels of CsgA, CsgD, and CsgG. In agreement with the results of CR-plate assays (Fig 1A), no CsgA was detected in the strains Δ*cbpA* Δ*dnaJ* and Δ*cbpA* Δ*djlA* Δ*dnAJ* as in the curli-negative strains Δ*csgA, ΔcsgD, ΔcsgG*, and Δ*rpoS* (Fig 3A and B). Likewise, a low level of CsgD was detected in the strains Δ*cbpA* Δ*dnaJ* and Δ*cbpA* Δ*djlA* Δ*dnaJ*. In addition, the cellular levels of CsgG in the strains Δ*cbpA* Δ*dnaJ* and Δ*cbpA* Δ*djlA* Δ*dnaJ* were lower than those in the wild-type strain. These data for the strains Δ*cbpA* Δ*dnaJ* and Δ*cbpA* Δ*djlA* Δ*dnaJ* are similar to those of the strain Δ*dnaK*. Our previous study indicated that expression of the *csgDEFG* operon was reduced in the strain Δ*dnaK* at the transcription level, which resulted in a reduction in the expression of the *csgBAC* operon (Sugimoto et al., 2018). Collectively, these results suggest that reduced expression of *csgDEFG* operon leads to the decreased expression of *csgBAC* operon in the strains Δ*cbpA* Δ*dnaJ* and Δ*cbpA* Δ*djlA* Δ*dnaJ*.

Next, we analyzed the cellular levels of RpoS, the stationary phase-specific sigma factor, in the BW25113 derivatives as it positively regulates the expression of the *csgDEFG* operon. A slight reduction in the RpoS level was observed in the strains Δ*dnaK, ΔcbpA ΔdnaJ*, and Δ*cbpA* Δ*djlA* Δ*dnaJ* (Fig 3). The decreased level of RpoS in the strain Δ*dnaK* may be due to accelerated proteolytic degradation by ClpXP (Rockabrand et al., 1998). Consistent with these results, the cellular activity of RpoS through the measurement of catalase activity was reduced in these mutant strains (Fig EV2), suggesting that a partially reduced RpoS level resulted in a decrease in the expression of the *csgDEFG* operon. However, the activity of RpoS in the strains Δ*dnaK, ΔcbpA ΔdnaJ*, and Δ*cbpA* Δ*djlA* Δ*dnaJ* was not low enough to shut the expression of the *csgDEFG* operon down. Therefore, other mechanisms may account for the drastic reduction of the cellular CsgD level in the strain Δ*cbpA* Δ*dnaJ* and Δ*cbpA* Δ*djlA* Δ*dnaJ*.

### Either DnaJ or CbpA is involved in folding of transcriptional regulator MlrA

To investigate whether either DnaJ or CbpA is important for the expression of CsgD (Fig 3), we tested the contribution of these JDPs to the folding of MlrA *in vitro*. MlrA was synthesized using an *in vitro* translation system (PURE System; Shimizu et al., 2001), in the presence or absence of complete and incomplete sets of DnaK/DnaJ/GrpE (KJE) or DnaK/CbpA/GrpE (KAE). *De novo* synthesized MlrA readily formed aggregates and a complete set of KJE or KAE assisted the folding of MlrA (Fig 4A and EV3A). Incomplete sets of DnaK/DnaJ (KJ), DnaJ/GrpE (JE) and DnaJ alone (J) also promoted the solubility of MlrA, indicating that the folding of MlrA strongly relied on DnaJ. It should be noted that CbpA can compensate for DnaJ in KJE-assisted folding of MlrA (Fig 4A and EV3A). In contrast, DnaK alone (K), DnaK/CbpA (KA), DnaK/GrpE (KE), CbpA/GrpE (AE), CbpA alone (A), and GrpE (E) showed no or only slight stimulation of the solubility of MlrA (Fig 4A and EV3A). These results indicate that either DnaJ or CbpA is required for the DnaK chaperone system to efficiently fold MlrA. In addition, these findings are in good agreement with those already described, indicating that either DnaJ or CbpA is indispensable for efficient curli production (Fig 1–3).

**Figure 4.**
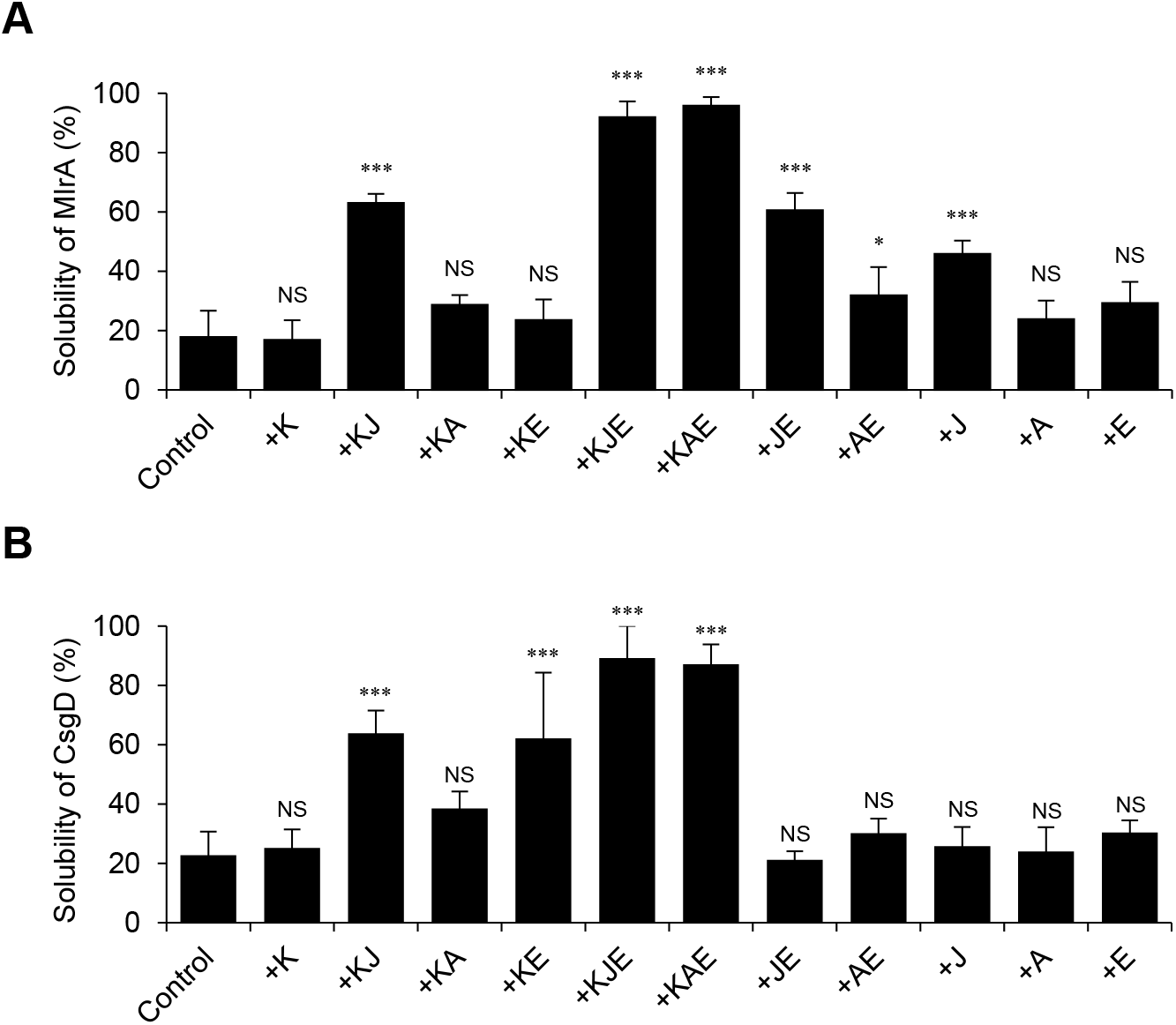
Effects of complete and incomplete DnaK chaperone systems on *de novo* folding of transcriptional regulators. (A, B) *De novo* folding of MlrA and CsgD was analyzed in the absence (Control) or presence of the indicated chaperone proteins using a cell-free translation system (PURE System). The solubilities (%) of MlrA (A) and CsgD (B) were calculated based on the band intensity of immunoblots. All experiments were repeated at least three times to ensure accuracy and averaged values with standard deviations were calculated. K, DnaK; KJ, DnaK/DnaJ; KA, DnaK/CbpA; KE, DnaK/GrpE; KJE, DnaK/DnaJ/GrpE; KAE, DnaK/CbpA/GrpE; JE, DnaJ/GrpE; AE, CbpA/GrpE; J, DnaJ; A, CbpA; E, GrpE. ***, P < 0.001. *, P < 0.05. NS, not significant.

Next, we examined whether *de novo* folding of CsgD required either DnaJ or CbpA, because its folding is assisted by the complete DnaK chaperone system (KJE) (Sugimoto et al., 2018). Our study revealed that CsgD formed aggregates in the absence of the chaperones and that KJE and KAE promoted CsgD folding (Fig 4B and EV3B). In contrast, DnaK alone did not support CsgD folding under the tested conditions (Fig 4B and EV3B). Interestingly, incomplete sets of the DnaK systems (KJ and KE) moderately enhanced the solubility of CsgD *in vitro* (Fig 4B and EV3B). DnaJ, CbpA, GrpE, and combinations of DnaJ and GrpE or CbpA and GrpE did not assist the folding of CsgD (Fig 4B and EV3B). These results indicate that these incomplete DnaK systems (KJ and KE) can act as molecular chaperones and that JDPs are dispensable at least for the folding of CsgD.

### Either DnaJ or CbpA is required for maintenance of CsgA in a translocation-competent state

Transport of CsgA across the cytoplasmic membrane is pivotal for curli biogenesis. In this process, DnaK acts on the transport precursor of CsgA to maintain its transport competent state via direct interaction with its N-terminal aggregation-prone signal peptide (Sugimoto et al., 2018). Firstly, we examined whether co-expression of CsgBAEFG was able to complement the defect of curli production in the strain BW2513 Δ*dnaJ* Δ*cbpA*, in which CsgA was not detected, as shown in Fig 3. Previously, we showed that introduction of the plasmid pCsgBAEFG was able to recover the production of curli in the BW25113 strains Δ*csgA, ΔcsgB, ΔcsgE*, ΔcsgF, and Δ*csgG*, confirming the plasmid was functional (Sugimoto et al., 2018). However, expression of CsgBAEFG did not restore the production of curli in Δ*dnaJ* Δ*cbpA* (Fig 5A), suggesting that secretion of CsgA was defective at the step of translocation from the cytoplasm to the periplasm or from the periplasm to the extracellular milieu.

**Figure 5.**
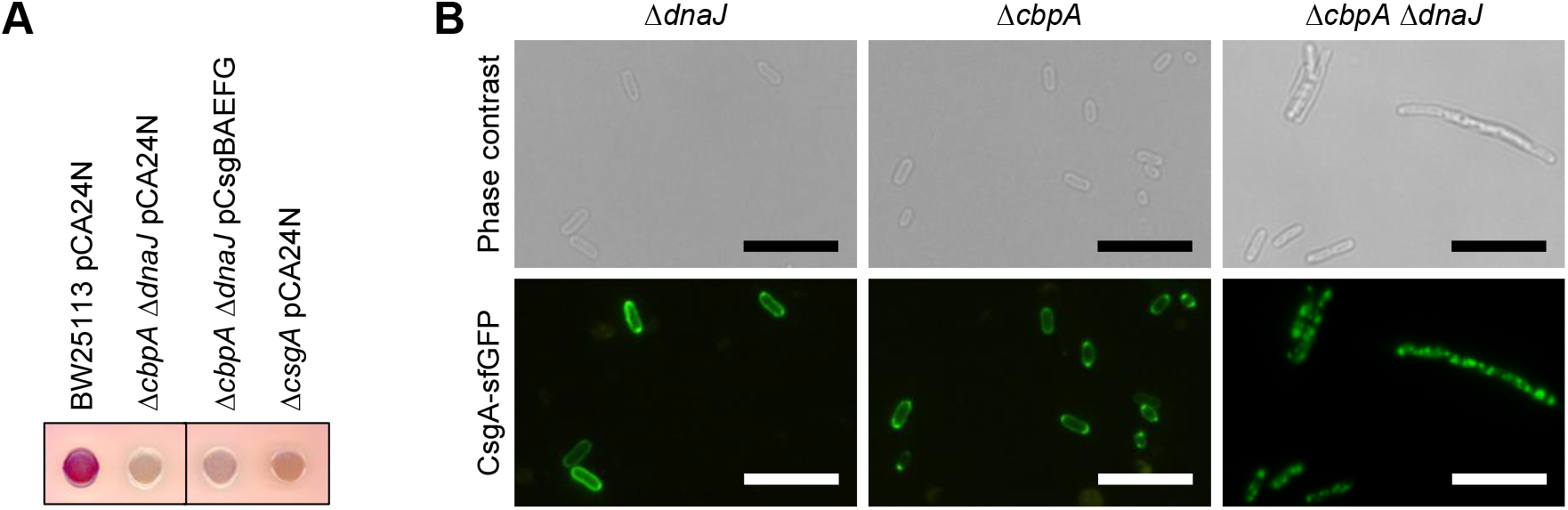
Effects of DnaJ and CbpA on translocation of CsgA-sfGFP. (A) Curli production in the indicated strains was examined on CR-YESCA plates supplemented with chloramphenicol as described in Fig 1B. The plates were incubated at 25°C for 2 days. (B) Translocation of CsgA across the cytoplasmic membrane was analyzed in BW25113 derivatives using a CsgA-sfGFP fusion protein (Sugimoto et al., 2018). *E. coli* cells were grown in LB medium supplemented with 100 μg/ml ampicillin. Scales, 10 μm.

Secondly, we investigated the requirements of JDPs for the translocation of CsgA from the cytoplasm to the periplasm using an *in vivo* visualization system (Sugimoto et al., 2018). Previously, we expressed a CsgA-sfGFP fusion protein in the BW25113 parental strain and its isogenic Δ*dnaK* strain. Subsequently, we observed that CsgA-sfGFP localized at the periplasm of the parental strain, whereas it formed aggregates in the cytoplasm of the Δ*dnaK* mutant (Sugimoto et al., 2018). In this study, we expressed the fusion protein in BW25113 derivative strains Δ*dnaJ, ΔcbpA*, and Δ*dnaJ* Δ*cbpA*. Fluorescence of sfGFP was observed at the periphery of the single null strains Δ*dnaJ* and Δ*cbpA*, indicating that CsgA-sfGFP was translocated to the periplasm. In contrast, it formed aggregates in the cytoplasm of the double knockout strain Δ*dnaJ* Δ*cbpA* (Fig 5B). These results suggest the requirement of either DnaJ or CbpA for translocation of CsgA from the cytoplasm to the periplasm.

Taken together, our results indicate that either DnaJ or CbpA is required for DnaK to assist the translocation of CsgA to the periplasm during curli biogenesis.

## DISCUSSION

Previously, we showed that DnaK is essential for curli biogenesis and biofilm formation via the maintenance of certain important proteins including RpoS, CsgD, and CsgA (Sugimoto et al., 2018). Here, we demonstrated that DnaJ and CbpA were essential for this process, while DjlA was not essential and that DnaJ and CbpA work redundantly in this process (Fig 1–3 and EV1). DnaK cooperates with DnaJ to assist in protein folding and to repair damaged proteins under harmful conditions, such as high temperature at 43°C, which causes denaturation and aggregation of numerous proteins (Fig 1C and EV1). The DnaK system efficiently prevents the aggregation of diverse thermolabile proteins, including 150–200 species, corresponding to 15–25% of detected proteins, under physiological heat stress conditions (Mogk et al., 1999). In addition, the disaggregation activities of the DnaK/CbpA/GrpE and DnaK/DjlA/GrpE systems were lower than that of the DnaK/DnaJ/GrpE system (Gur et al. 2004), suggesting that only DnaJ is essential for survival under severe stress conditions (e.g., heat stress at ≥43°C). Our model is consistent with the current body of research in several respects: (1) In curli biogenesis, only a subset of proteins (minimally RpoS, CsgD, and CsgA) require DnaK for their correct folding or export (Sugimoto et al., 2018). Our study also suggests that efficient folding of MlrA needs the DnaK system including DnaK, GrpE, and either DnaJ or CbpA (Fig 3–5). (2) DnaK mutants (DnaK^YND^, DnaK^K155D^, and DnaK^D393A^) with reduced basal activity were functional in the curli production but not in survival at high temperature of 43°C (Fig 2). (3) *E. coli* produces curli during the stationary phase of growth, in which expression of CbpA is induced via RpoS (Fig 3A). Therefore, the contribution of CbpA to the promotion of the DnaK function may be emphasized during curli biogenesis. This growth phase-specific selection of JDPs is a reasonable strategy for improving bacterial survival and fitness under nutrient-depleted conditions (e.g., during stationary phase) and adaptation during host colonization. (4) DjlA is an inner membrane anchoring JDP in *E. coli* (Clarke et al., 1996). Therefore, DjlA may be involved in the quality control of membrane proteins and trafficking of exported proteins (Kelley and Georgopoulos, 1997). However, DjlA is dispensable for maintenance of proteins associated with curli production.

Based on these observations, we propose a hierarchical model whereby Hsp70 chaperone activities regulate proteostasis in distinct cellular functions (Fig 6). When a large amount/variety of proteins in the cell are injured by severe stress conditions (e.g., heat stress at ≥43°C), full specification of the DnaK system (DnaK/DnaJ/GrpE) prevents their aggregation and the repair of toxic protein aggregates. In contrast, under certain conditions (e.g., curli biogenesis), a moderately active DnaK system (DnaK/CbpA/GrpE) fulfills its chaperone function by handling only a subset of proteins. Although DnaK/DjlA/GrpE and DnaK alone did not support the production of curli, they may function weakly by holding substrate proteins (Evans et al., 2011), which might be associated with a specific phenotype such as colanic acid production (Kelley and Georgopoulos, 1997) and other unknown cellular functions. This model is also supported by the results of DnaK mutants in which DnaK^YND^, DnaK^K155D^, and DnaK^D393A^ restored the curli production in the Δ*dnaK* strain despite the failure to support the growth at high temperature (Fig 2). In addition, neither DnaK^K70A^ which possesses a defective ATPase activity nor DnaK^V436F^ which retains decreased substrate affinity was able to rescue the thermosensitivity as well as the deficiency in curli production of the Δ*dnaK* strain (Sugimoto et al., 2018).

**Figure 6.**
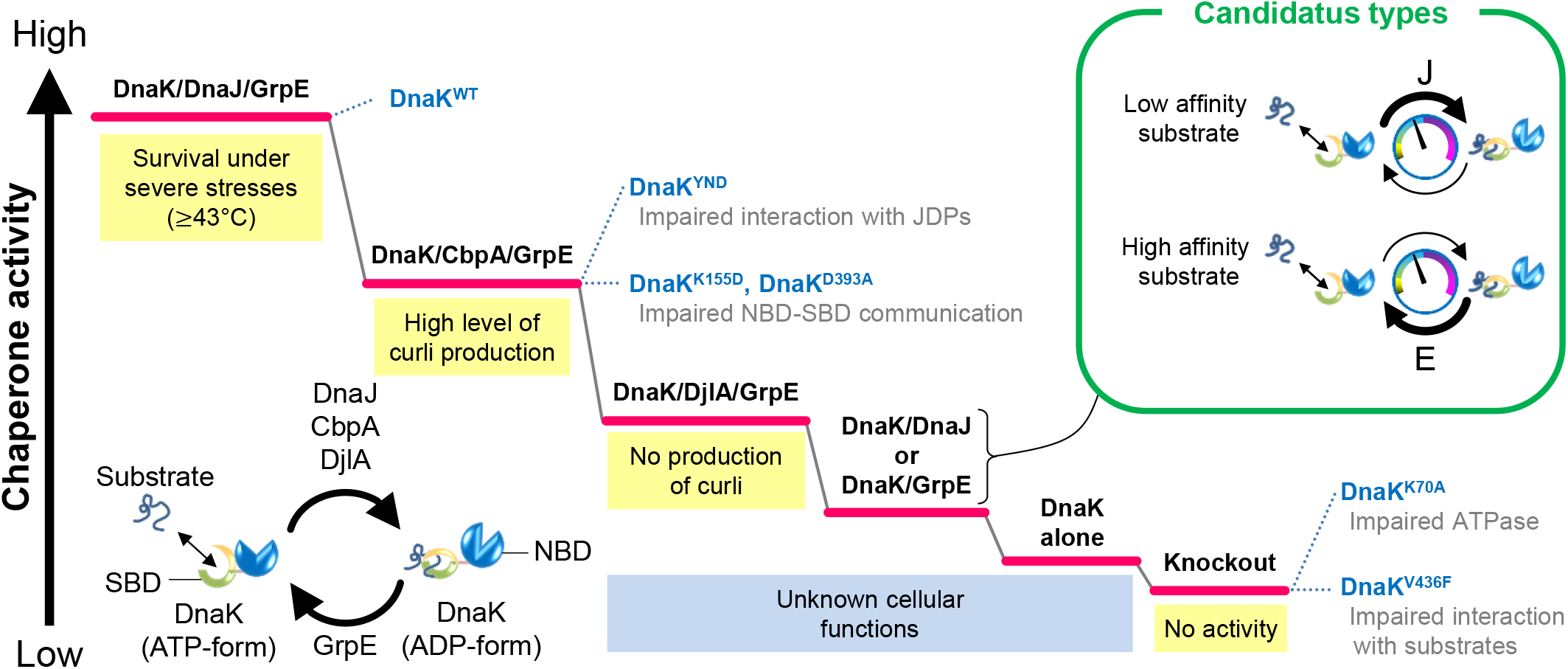
A hierarchical model of DnaK chaperone activities. The proposed model for DnaK chaperone activities showing that the most powerful DnaK/DnaJ/GrpE system is essential for survival under severe stress conditions (e.g., high temperature) for dealing with a wide variety of substrate proteins. The middle active DnaK/CbpA/GrpE system (Gur et al. 2004) can perform specific cellular functions (e.g., curli biogenesis). The modest active DnaK/DjlA/GrpE system is not able to support curli production but may work for unknown cellular activities. DnaK alone engages in holding substrate proteins, but its cellular functions remain elusive. Respective DnaK mutants with hierarchical activities are indicated based on the results presented in this study and our previous one (Sugimoto et al., 2018).

This study focused on the requirement of JDPs in the production of bacterial amyloid fibers. In addition to JDPs, DnaK cooperates with GrpE, a well-conserved bacterial NEF. We were also interested in the requirement of GrpE in curli biogenesis. However, deletion of the *grpE* gene from the genome of BW25113 was difficult because of its lethality in *E. coli* (Ang and Georgopoulos, 1989). Genetic analysis using an available *grpE* null strain derived from *E. coli* strain C600 (Ang and Georgopoulos, 1989; Sugimoto et al., 2008) suggested that GrpE may be dispensable for curli production (Sugimoto et al., unpublished). Further careful study is needed to clarify this observation.

Our data provide insight into the evolution of molecular chaperones and proteostasis. It has been previously suggested that Hsp70 chaperone systems co-evolved with the proteome to regulate the physiological state of the cell (Powers and Balch, 2013). The number of Hsp70s increases roughly linearly as the size of the genome increases (Powers and Balch, 2013). In addition, the number of JDPs is often much higher than that of Hsp70s in almost all life forms (Kampinga and Craig, 2010). Moreover, in contrast to Hsp70s, JDPs show a greater degree of sequence and structural divergence. These insights imply that they may play a major role in driving the multi-functionality of Hsp70 chaperone systems (Craig and Marszalek, 2017). In contrast, the single set of DnaK-DnaJ-GrpE is well conserved in Gram-positive bacteria, such as *Staphylococcus aureus* (Table EV1) (Warnecke, 2012). Generally, the genome sizes of Gram-positive bacteria are smaller than those of Gram-negative bacteria. Therefore, smaller numbers of Hsp70s and JDPs in Gram-positive bacteria may be anticipated, as the proteome size is also small, minimizing the work of the Hsp70 system. In the cases of microorganisms with extremely small genomes, such as Candidatus *Hodgkinia cicadicola* and Candidatus *Carsonella ruddii*, only either DnaJ or GrpE is present, and there is a single Hsp70/DnaK (Table EV1). How do such incomplete DnaK systems (so called proto-DnaK systems) work in maintaining proteostasis? These organisms possess quite small numbers of proteins; therefore, the roles of the DnaK systems must be minimal. In this situation, activity of proto-DnaK systems (DnaK/DnaJ and DnaK/GrpE) may be sufficient for regulating a limited proteome to aid survival of these microorganisms (Fig 6). This notion is consistent with our results that incomplete sets of the *E. coli* DnaK chaperone system (DnaK/DnaJ and DnaK/GrpE) can contribute to folding of certain proteins (e.g., MlrA and CsgD) (Fig 4B and EV3B). Whether other cellular processes require either full-specification DnaK systems or only proto-DnaK systems remains unclear; some proteins may require DnaK and either DnaJ or GrpE for their folding. It is expected that proteins with lower affinity with DnaK need only DnaJ, since DnaK alone is not able to capture them and these proteins can be released spontaneously from DnaK despite the absence of GrpE. In contrast, folding of proteins with higher affinity with DnaK may depend on GrpE because, although DnaK alone is able to bind them, the release of proteins tightly bound to DnaK requires GrpE. The lower affinity proteins may be expressed predominantly in Candidatus bacteria that possess the DnaK/DnaJ system, whereas the higher affinity proteins may be expressed in Candidatus bacteria that possess the DnaK-/GrpE system. These insights into diversification and evolution of the Hsp70 chaperone system, in combination with our data, imply that primitive organisms may use proto-DnaK systems to manage their small proteomes.

## METHODS AND MATERIALS

### Bacterial strains

The *E. coli* strains used in this study are listed in Table EV2. All strains were cultivated in LB medium (1% tryptone, 0.5% yeast extract, 0.5% NaCl) or YESCA medium (1% casamino acid, 0.1% yeast extract). When appropriate, the medium was supplemented with 30 μg/ml chloramphenicol, 50 μg/ml kanamycin, or 100 μg/ml ampicillin.

### Construction of *E. coli* null mutant strains

The JDP-null mutant strains of BW25113 (Table EV2) were constructed by the one-step method for inactivation of chromosomal genes (Datsenko and Wanner, 2000; Baba et al., 2006). The plasmids and primers used for gene knockout are listed in Tables EV2 and EV3, respectively.

### Construction of plasmids

The ASKA clone plasmids (pASKA-DnaJ, pASKA-CbpA, pASKA-DjlA, pASKA-CsgD, and pASKA-MlrA) were provided by the National Institute of Genetics (Shizuoka, Japan). For construction of pDjlA^ΔTM^ (Table EV2), the DNA encoding the transmembrane domain was deleted by inverse PCR using KOD Plus Neo DNA polymerase (Toyobo, Osaka, Japan), pASKA-DjlA as a template, and a primer set DjlA-deltaTM-F and DjlA-deltaTM-R (Table EV3).

For construction of plasmids expressing DnaK mutants (DnaK^YND^, DnaK^K155D^, and DnaK^D393A^), site-directed mutagenesis was performed by inverse PCR using KOD Plus Neo DNA polymerase (Toyobo, Osaka, Japan), pDnaK^WT^ as a template, and the following primer sets: dnaK-YND-F/dnaK-YND-R, dnaK-K155D-F/dnaK-K155D-R, and dnaK-D393A-F/dnaK-D393A-R. The resultant plasmids were termed pDnaK^YND^, pDnaK^K155D^, and pDnaK^D393A^, respectively (Table EV2).

The plasmids were analyzed by DNA sequencing (Eurofins Genomics, Tokyo, Japan). Primers used in this study were synthesized by Thermo Fisher and are summarized in Table EV3.

### Protein purification

Recombinant DnaK, DnaJ, and GrpE were purified as described previously (Niwa et al., 2012). CbpA was expressed in *E. coli* BL21(DE3). Cells harboring pCU60 were grown at 30°C in 2× YT medium containing 100 μg/ml ampicillin, and expression of CbpA was induced by adding IPTG (1 mM) and incubating at 30°C for 3 h. Cells from 2-L culture were harvested by centrifugation and resuspended in 50 ml buffer A [10 mM Tris-HCl (pH 8.0), 1 mM DTT, 10% glycerol] supplemented with a protease inhibitor cocktail. After sonication on ice, cell lysates were centrifuged at 12,000 ×*g* for 60 min at 4°C, and the supernatant was loaded onto a 5-ml bed volume of HiTrap Heparin column (GE Healthcare, Pittsburgh, PA, USA) pre-equilibrated with buffer A. CbpA was eluted using a 0– 1,000 mM NaCl gradient in buffer A. Each fraction containing CbpA was pooled and further purified by chromatography using a HiTrap Q column (GE Healthcare) and a 0–1,000 mM NaCl gradient in buffer A. Purified CbpA was confirmed by LC-MS/MS and quantified using a Bradford Assay Kit.

### Antibodies

Rabbit anti-DnaJ and rabbit anti-RpoH antisera were gifted by Dr. B. Bukau (Gamer et al., 1992). The other antibodies were prepared as previously reported (Arita-Morioka et al., 2018; Sugimoto et al., 2018).

### Congo Red (CR)-binding assay

Curli formation was assayed at 25°C on CR-containing YESCA (1% casamino acid, 0.1% yeast extract, 2% agar) plates as previously reported (Arita-Morioka et al., 2018; Sugimoto et al., 2018). When needed, 30 μg/ml chloramphenicol was added to supplement select transformants.

### Thermosensitivity assay

*E. coli* BW25113 derivative cells were grown at 30°C in LB medium overnight. Overnight cultures were serially diluted 10-fold, and 5 μl of these dilutions were spotted onto LB agar plates. If required, 30 μg/ml chloramphenicol was added to supplement select transformants. These plates were incubated at 30°C or 43°C for 24 h.

### Immunoblotting

For detection of CsgA, CsgD, CsgG, RpoS, RpoD, RpoH, DnaK, DnaJ, CbpA, and His-tagged MlrA, immunoblotting was performed as previously reported (Arita-Morioka et al., 2018; Sugimoto et al., 2018). After SDS-PAGE, proteins were transferred to polyvinylidene difluoride membranes using the iBlot 2 dry blotting system (Thermo Fisher) following the manufacturer’s instructions. Membranes were blocked with blocking solution [1–5% skimmed milk, Tris-buffered saline containing 0.1% (v/v) Tween 20 (TBS-T)] for at least 1 h at 25°C or overnight at 4°C. After gentle washing with TBS-T, the membrane was incubated with appropriate primary antibodies for at least 1 h at 25°C or overnight at 4°C. Membranes were subsequently probed with appropriate secondary HRP-conjugated antibodies for 1 h at 25°C or overnight at 4°C. After washing the membrane three times with TBS-T, signals were detected using the ECL Prime Western Blotting Detection Reagent (GE Healthcare) and an ImageQuant LAS-4000 system (GE Healthcare). When required, signal intensities were quantified with ImageQuant TL software version 7.0 (GE Healthcare).

Primary antibodies were diluted into CanGet Signal 1 (Toyobo) as follows: anti-CsgA (1/1,000), anti-CsgD (1/200), anti-CsgG (1/5,000–1/1,000), anti-RpoS (1/10,000–1/1,000), anti-RpoD (1/10,000–1/1,000), anti-RpoH (1/2,000), anti-DnaK (1/10,000), anti-DnaJ (1/10,000), anti-CbpA (1/10,000), and anti-His (1/10,000). HRP-conjugated goat anti-rabbit IgG and HRP-conjugated goat anti-mouse IgG secondary antibodies were diluted 1/50,000 and 1/10,000, respectively, in CanGet Signal 2 (Toyobo).

For detection of CsgA monomers, curli fibers were depolymerized into subunits by treatment with hexafluoroisopropanol (HFIP) before SDS-PAGE (Sugimoto et al. 2018). Bacterial cells (1 mg) were suspended in 10 μl STE buffer [10 mM Tris-HCl (pH 8.0), 100 mM NaCl, 2 mM EDTA] and mixed well with 50 μl HFIP. After brief sonication, samples were vacuum dried using a SpeedVac vacuum concentrator (Thermo Fisher) at 45°C for more than 30 min. The HFIP-treatment was repeated. The dried materials were dissolved in 40 μl 8 M urea solution. After brief sonication in a water bath for 5 min at room temperature, the solutions were mixed with an equal volume of 2× SDS sample buffer [150 mM Tris-HCl (pH 6.8), 4% SDS, 20% glycerol, 10% 2-mercaptoethanol]. Proteins were separated by 15% SDS-PAGE.

### *In vitro* protein folding assay

*De novo* folding of CsgD and MlrA was analyzed using the PURE System (Shimizu et al., 2001) as previously reported (Niwa et al., 2012; Sugimoto et al., 2018). The *csgD* gene was amplified from the CsgD-expression plasmid pASKA-CsgD (Table EV2) by PCR using KOD Plus DNA polymerase v. 2 (Toyobo) and the primer set Pure-Niwa-F and Pure-CsgD-R (Table EV3). The *mlrA* gene was amplified from the MlrA-expression plasmid pASKA-MlrA (Table EV2) by PCR using Phusion High-Fidelity DNA polymerase (New England Biolabs, Tokyo, Japan) and the primer set Pure-Niwa-F and Pure-Niwa-R (Table EV3). The amplified DNA fragments were purified and incubated with PURE*frex* solution (GeneFrontier Corp., Chiba, Japan) at 37°C for 3–4 h according to the manufacturer’s instructions. When required, reaction mixtures (20–40 μl) were supplemented with DnaK (5 μM), DnaJ (1 μM), CbpA (1 μM), and/or GrpE (1 μM). After incubation, aliquots (10–20 μl) of the solution were obtained as the total fractions and were centrifuged at 20,000 *×g* for 30 min at 4°C to separate the soluble and insoluble fractions. The equivalent volumes of the total, soluble, and insoluble fractions were mixed with 2× SDS sample buffer. After boiling at 95°C for 5 min, proteins were resolved by 15% SDS-PAGE and stained with CBB. For detection of CsgD and MlrA, immunoblotting was performed as described above.

### Fluorescence microscopy

Transport and aggregation of CsgA-sfGFP was observed in *E. coli* as previously reported (Sugimoto et al., 2018) with a slight modification. *E. coli* expressing CsgA-sfGFP were grown in LB medium supplemented with 100 μg/ml ampicillin at 30°C overnight. A small aliquot of the overnight cultures was placed on a slide and covered with a coverslip. Fluorescence of sfGFP was observed under a fluorescence microscope (Nikon, Tokyo, Japan) equipped with B2 (excitation filter, 450–490 nm; barrier filter, 520 nm) and G2A (excitation filter, 510–560 nm; barrier filter, 590 nm) filters. In this study, no arabinose was supplemented into the media, because leaky expression from the plasmid was sufficient to visualize fluorescence.

### Statistical analysis

One-way ANOVA with Dunnett’s post hoc test was used to determine whether any of the groups exhibited a statistically significant difference in the solubility of MlrA and CsgD analyzed by the PURE System. All experiments were performed at least three times. For all analyses, *P* < 0.05 was considered statistically significant.

## ACKNOWLEDGEMENTS

We acknowledge Dr. B. Bukau for providing the anti-DnaJ and anti-RpoH antibodies, Dr. Y. Ueguchi for gifting pCU60, Dr. N. Tani for LC-MS/MS analysis, National BioResource Project (NBRP, Japan) for the Keio collection and ASKA clone. We also thank A. Terao, N. Toda, N. Fukuda, D. Fujioka, and H. Iso for experimental assistance and other members in Department of Bacteriology, Jikei University School of Medicine for stimulating discussions. We are grateful to Dr. T Kanamori for supporting the analysis using the PURE System.

This work was partially supported by a Grant-in-Aid for Young Scientists (A) to S.S. from JSPS (15H05619), a Grant-in-Aid for Fund for the Promotion of Joint International Research [Fostering Joint International Research (A)] to S.S. from JSPS (18KK0429), a grant to Y.M. from the MEXT-Supported Program for the Strategic Research Foundation at Private Universities, 2012–2016, the Joint Usage/Research Center for Developmental Medicine, IMEG, Kumamoto University, The Uehara Memorial Foundation to S.S., and by JST ERATO Grant Number JPMJER1502.

## AUTHOR CONTRIBUTIONS

S.S., K.Y., and T.O. planned the project. S.S. designed the experiments and developed the assay. S.S. and T.N. purified proteins. S.S. performed the experiments and analyzed the data. Y.K. and Y.M. supported the project. S.S. and K.Y. wrote the paper with input from all co-authors.

## EXPANDED VIEW LEGENDS

**Figure EV1. Effects of JDP deletion on thermotolerance of *E. coli* strain BW25113**

(A) Thermosensitivity of the indicated strains was assayed on LB agar plates. Ten-fold serial dilutions of the overnight cultures were spotted on the plates and incubated at 30°C or 43°C for 24 h. (B) BW25113 and its isogenic Δ*cbpA* Δ*dnaJ* Δ*dnaJ* strains were transformed with pCA24N (empty vector) or the indicated JDP-expression plasmids. The strains were grown at 30°C or 43°C for 24 h on LB agar plates supplemented with chloramphenicol.

**Figure EV2. Activity of RpoS in BW25113 derivative strains**

Relative RpoS activity was assayed by measuring catalase activity as previously reported (Sugimoto et al., 2018). All experiments were repeated at least three times to ensure accuracy and averaged values with standard deviations were calculated. The activity of the BW25113 parental strain was defined as 100%.

**Figure EV3. Effects of complete and incomplete DnaK chaperone systems on de novo folding of transcriptional regulators MlrA and CsgD**

(A, B) De novo folding of MlrA and CsgD was analyzed in the absence (Control) or presence of the indicated chaperone proteins using a cell-free translation system (PURE System). His-tagged MlrA (A) and CsgD (B) were detected using anti-His antibody and anti-CsgD antibody, respectively. All experiments were repeated at least three times to ensure accuracy and representative images are shown. K, DnaK; KJ, DnaK/DnaJ; KA, DnaK/CbpA; KE, DnaK/GrpE; KJE, DnaK/DnaJ/GrpE; KAE, DnaK/CbpA/GrpE; JE, DnaJ/GrpE; AE, CbpA/GrpE; J, DnaJ; A, CbpA; E, GrpE.

